# Genetic reagents for making split-GAL4 lines in Drosophila

**DOI:** 10.1101/197509

**Authors:** Heather Dionne, Karen L. Hibbard, Amanda Cavallaro, Jui-Chun Kao, Gerald M. Rubin

## Abstract

The ability to reproducibly target expression of transgenes to small, defined subsets of cells is a key experimental tool for understanding many biological processes. The Drosophila nervous system contains thousands of distinct cell types and it has generally not been possible to limit expression to one or a few cell types when using a single segment of genomic DNA as an enhancer to drive expression. Intersectional methods, in which expression of the transgene only occurs where two different enhancers overlap in their expression patterns, can be used to achieve the desired specificity. This report describes a set of over 2,800 transgenic lines for use with the split-GAL4 intersectional method.

## Introduction

We previously reported the expression patterns of over 7,000 GAL4 lines in the adult nervous system of *Drosophila melanogaster* (Pfeiffer *et al*. 2008; Jenett *et al*. 2012; www.janelia.org/gal4-gen1). In these lines (referred to below as generation 1 GAL4 lines), expression of the transcription factor GAL4 is driven by a 2-3 kb segment of genomic DNA that contains one or more transcriptional enhancer sequences. While many lines show expression in a small fraction (less than 1%) of neurons in the adult brain, only about one in a thousand appears to drive expression in a single cell type (Jenett *et al*. 2012). To achieve greater specificity, we turned to the split-GAL4 intersectional method (Luan *et al*. 2006) using the optimized vectors described in Pfeiffer *et al.* (2010). In this method, individual enhancers drive the expression of either GAL4’s DNA-binding domain (DBD) or an activation domain (AD) joined to a leucine-zipper dimerization domain. When expressed individually, each half is insufficient to activate transcription of a UAS reporter. When both AD and DBD are present in the same cell, they combine to make a functional transcription factor that can bind to tandem arrays of GAL4’s cognate UAS DNA sequence and activate transcription.

Here we describe a set of such transgenic “hemidriver” lines expressing either the p65 AD or GAL4 DBD domain under the control of an enhancer from the collection described in Jenett *et al*. (2012). These lines have been used successfully to make comprehensive sets of lines that each show expression in one or a few cell types in a particular brain area, for example: the lamina of the optic lobe (Tuthill *et al*. 2013), mushroom body intrinsic and extrinsic neurons (Aso *et al*. 2014), and lobula columnar neurons (Wu *et al*. 2016). We discuss a sample screening protocol and the typical results we obtain as a practical guide for those who want to use the hemidriver lines to develop split-GAL4 lines specific for additional cell types.

## Results and Discussion

### Construction of hemidriver lines

Lines were constructed essentially as described by Pfeiffer *et al.* (2010) using entry clones generated as described in Pfeiffer *et al*. (2008). To transfer the enhancer regions to split-GAL4 destination vectors, approximately 50 ng of each entry clone was used in Gateway reactions with LR clonase (Thermo Fisher) and either pBPZpGAL4DBDUw or pBPp65ADZpUw (Pfeiffer *et al*. 2010; available from Addgene, plasmids 26233 and 26234, respectively). The p65 AD replaces the native GAL4 AD in pBPp65ADZpUw, which results in stronger transcriptional activation and insensitivity to inactivation by GAL80. BPp65ADZp and BPZpGDBD, hemidriver constructs that lack an enhancer fragment, were constructed by substituting the GAL4 coding sequence in pBPGAL4U (Pfeiffer *et al*. 2008) with the split-GAL4 coding sequences from pBPp65ADZpUw and pBPZpGAL4DBDUw, respectively, using *Kpn*I and *Hind*III sites. DNA for injections was prepared from 5 ml overnight cultures using a QIAprep kit (Qiagen) and verified by *EcoR*I restriction digestion. The HL9, Tdc2, TH and Trh hemidrivers are described in Aso *et al*. (2014).

The DBD hemidrivers were inserted using phiC31 site specific integrase into the attP2 (3L) landing site (Groth *et al*. 2004) and the AD hemidrivers were inserted in the attP40 (2L) landing site (Markstein *et al.* 2008). The injections to generate the transformants were performed by Genetic Services Inc. (Cambridge, MA). Newly generated transformants were processed through a series of genetic crosses to remove the integrase source and to establish a homozygous stock, as diagrammed in Supplemental Figure 1 of Pfeiffer *et al.* 2008. A subset of the DBD hemidrivers were balanced to facilitate further stock construction and efficient intersectional screening using the stock pJFRC200-10XUAS-IVS-myr::smGFP-HA in attP18, pJFRC216-13XLexAop2-IVS-myr::smGFP-V5 in su(Hw)attP8; wg^*Sp-^1^*^*/CyO*; *TM2/TM6B* (Nern *et al*. 2015). Some of the AD hemidrivers were balanced by a similar cross scheme utilizing one of the following stocks: *w; Amos*^*Tft*^*/CyO*; +*w; wg*^*Sp-^1^*^*/CyO; MKRS/TM6B, or w; Kr*^*If-^1^*^*/CyO,2xTb-RFP; MKRS/TM6B*. Approximately 90% of the hemidriver lines could be maintained as homozygous viable stocks. Those lines that were homozygous lethal, sterile, or sickly were maintained over a balancer. We believe that in most or all cases the lethality or sterility was due to background mutations in the stocks. We backcrossed ˜100 lethal or sterile AD lines for three generations to a *w*^*1118*^ stock and were able to establish homozygous stocks in 90 percent of these cases. The hemidriver lines shown in Table 1 have been deposited in the Bloomington Drosophila Stock Center (http://flystocks.bio.indiana.edu).

**Table 1.**
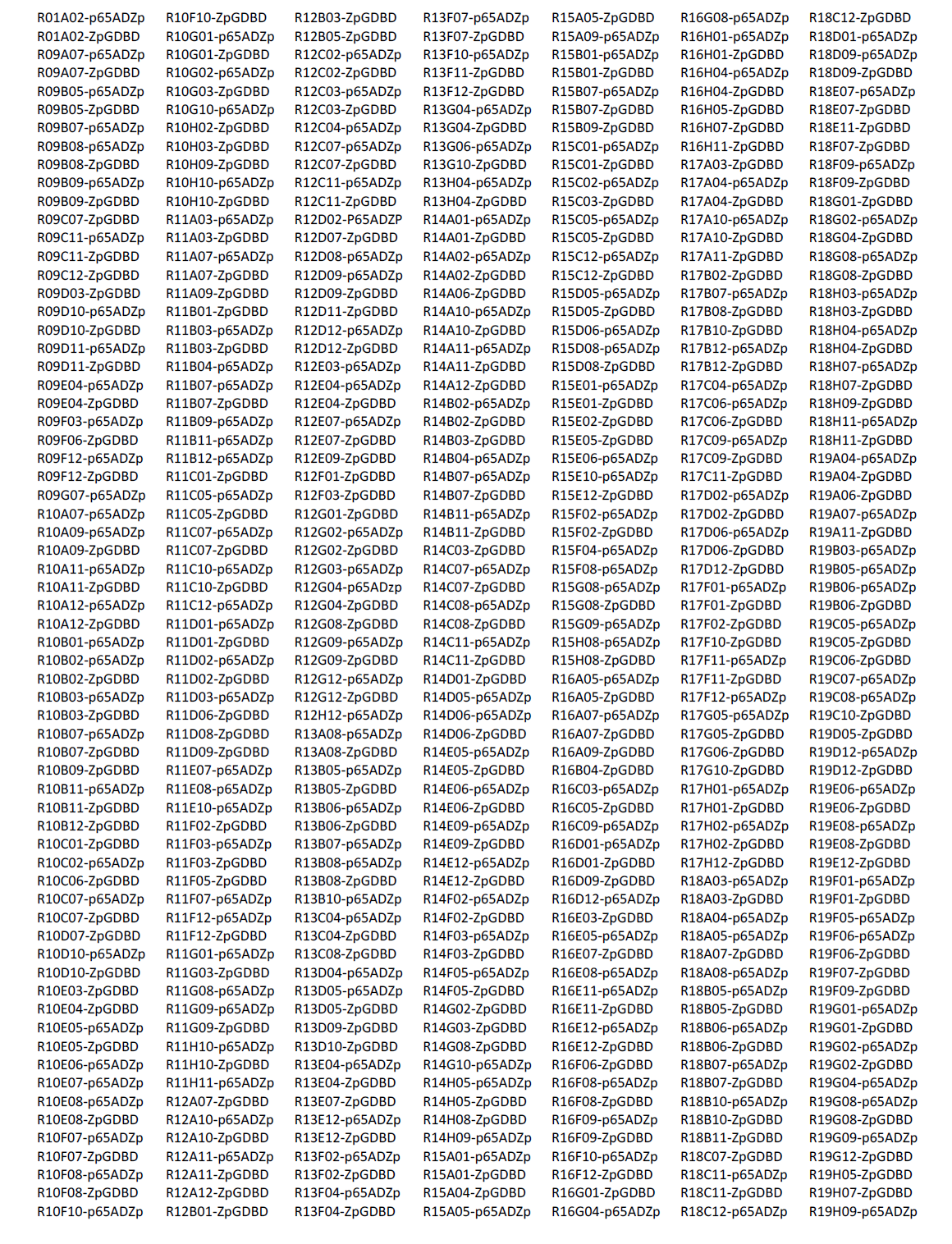
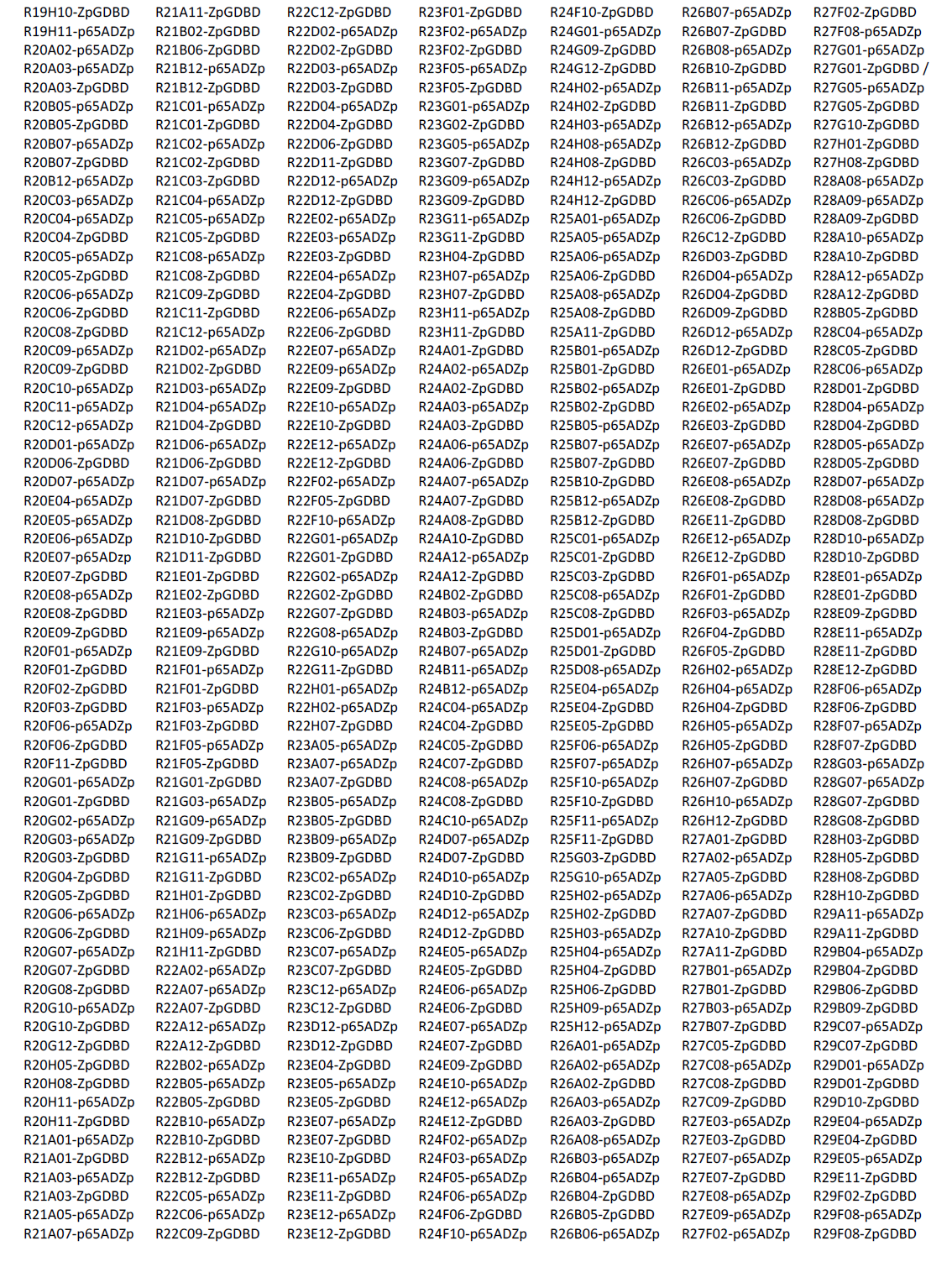
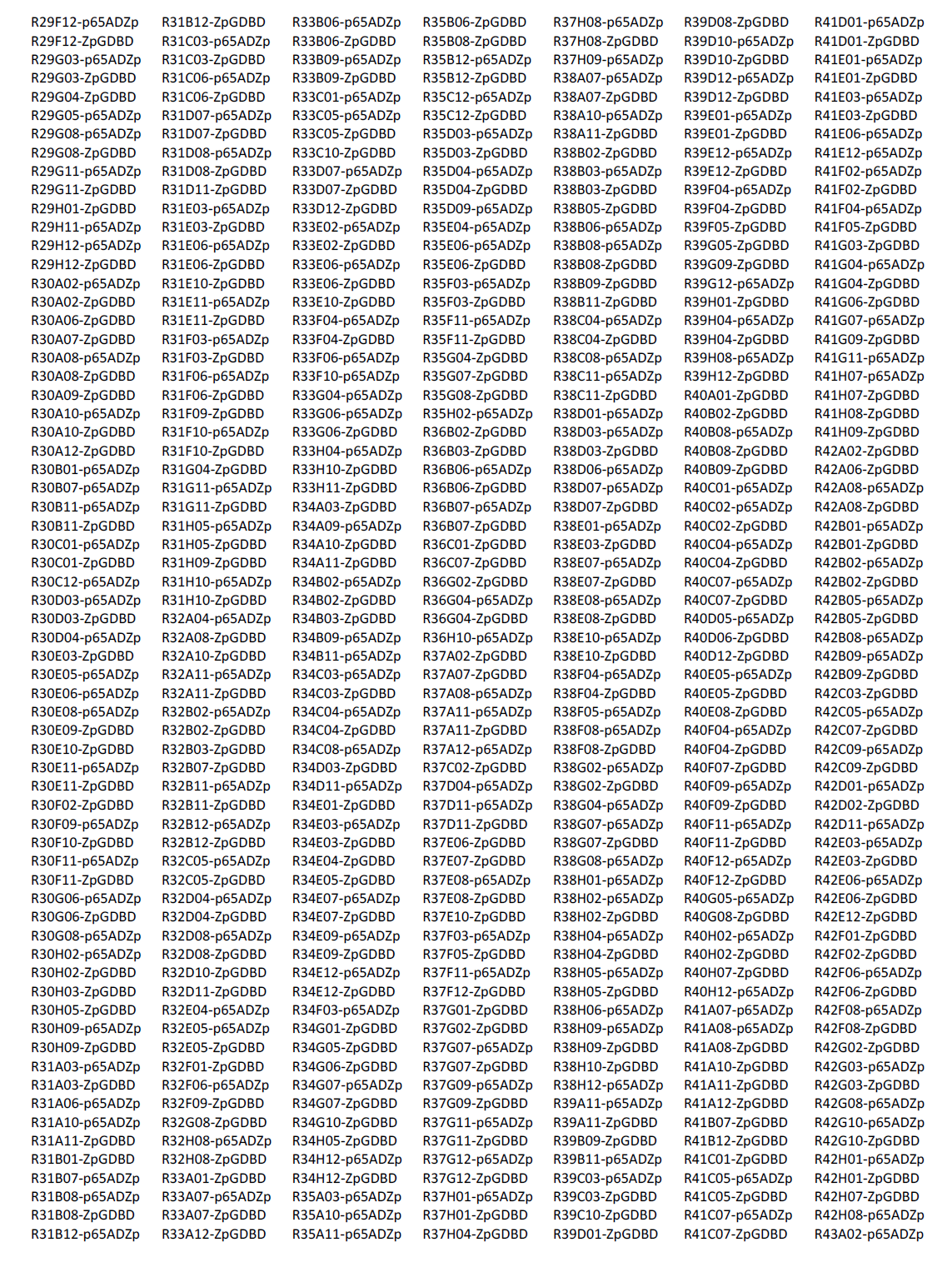
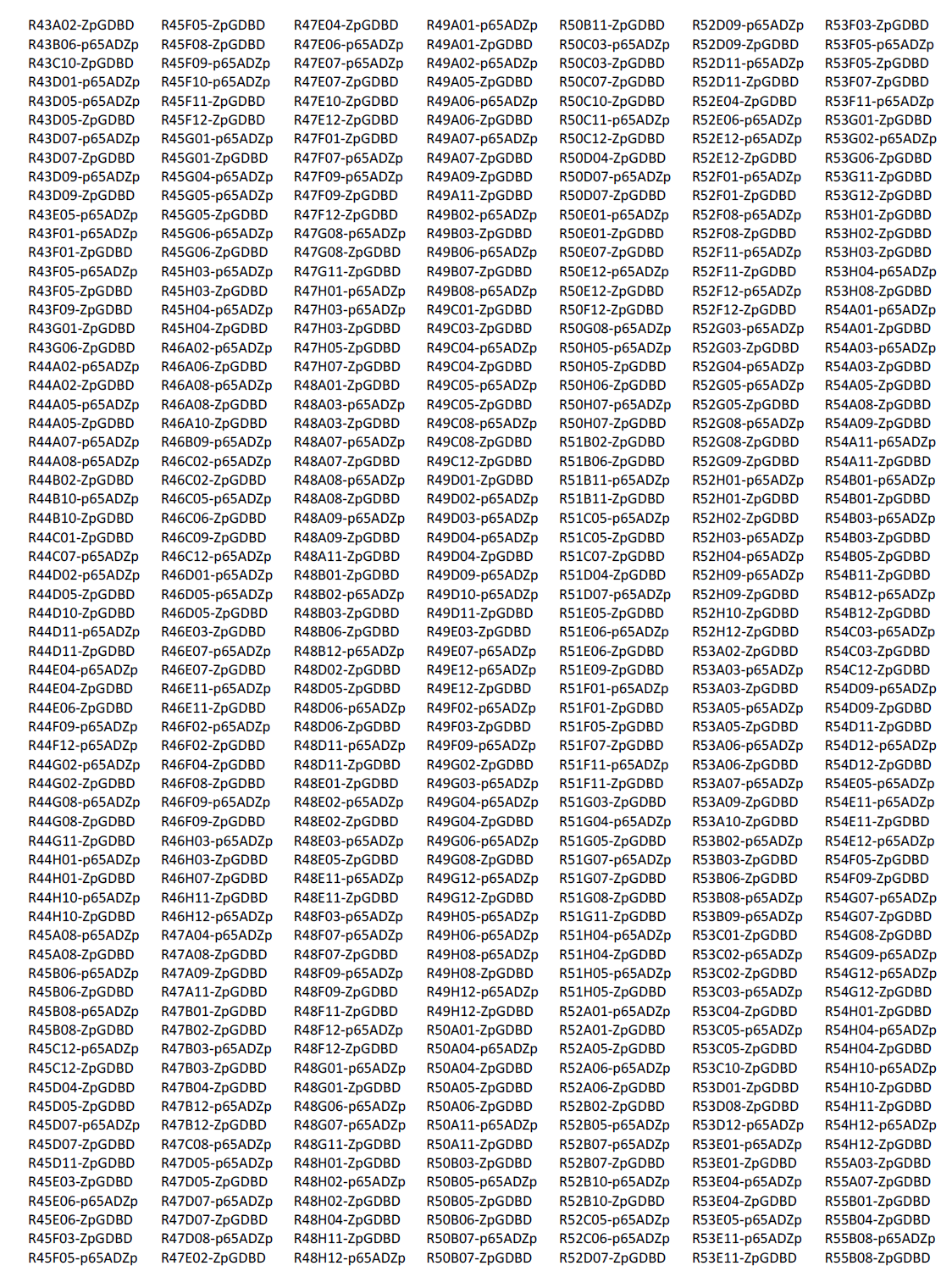
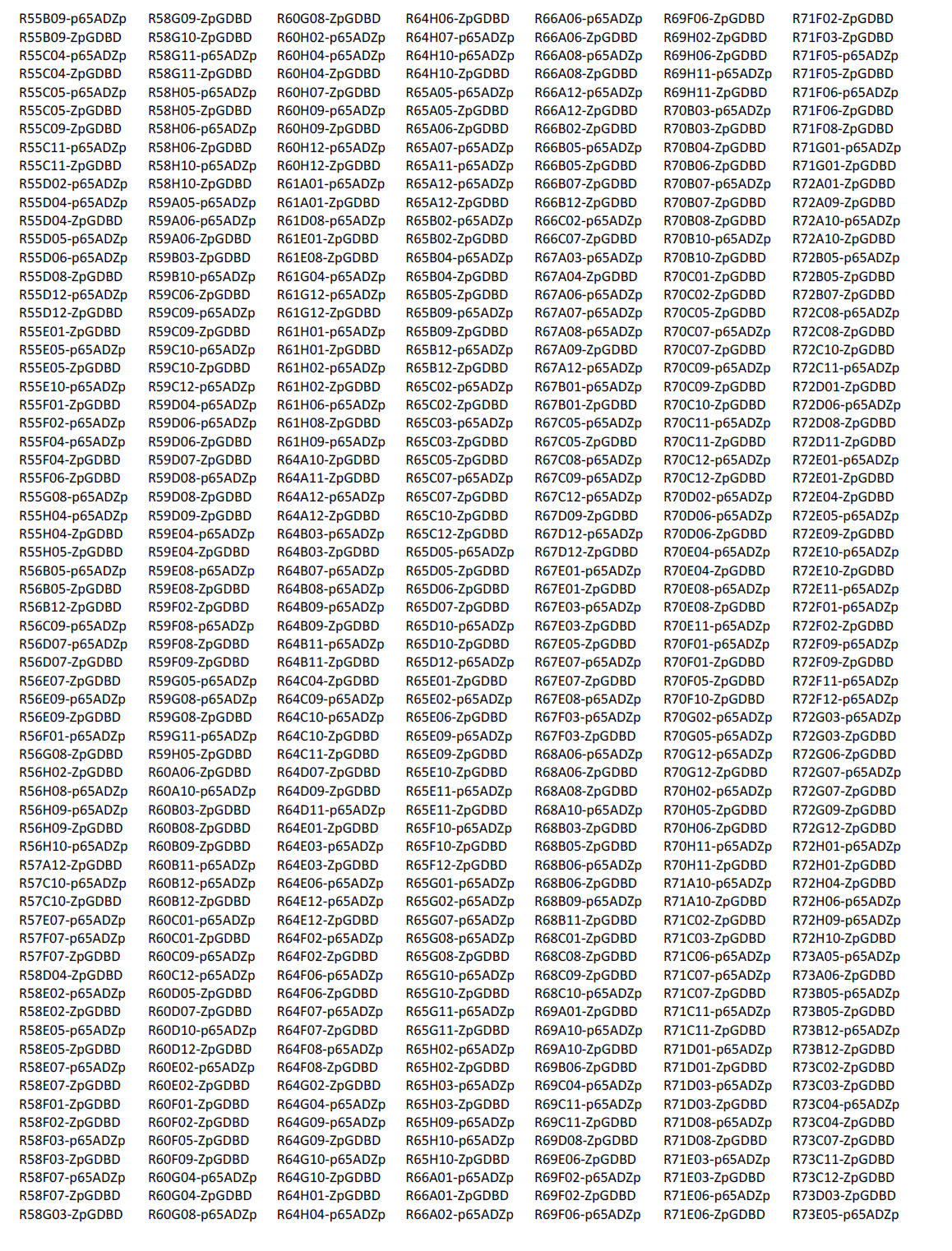
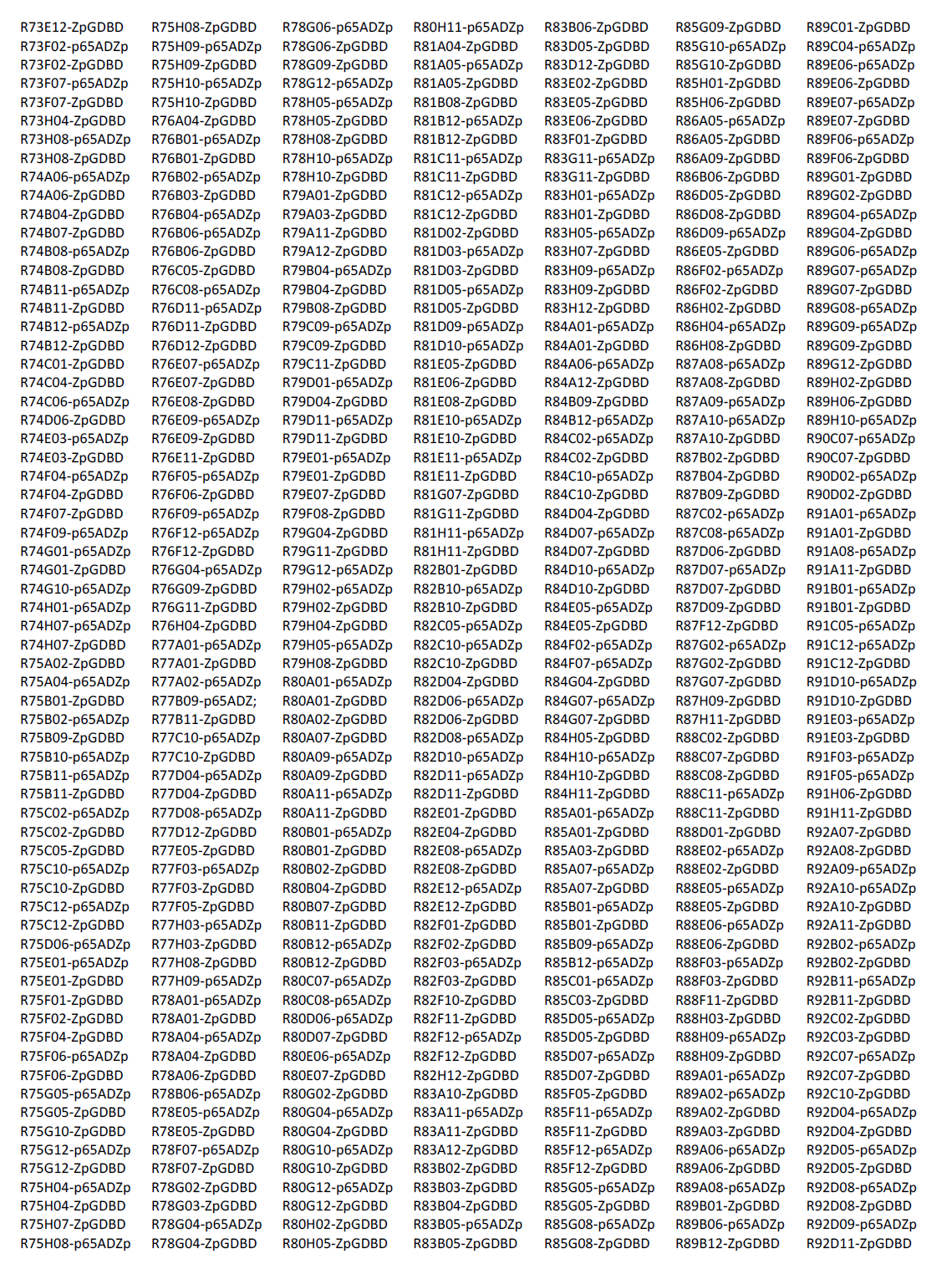
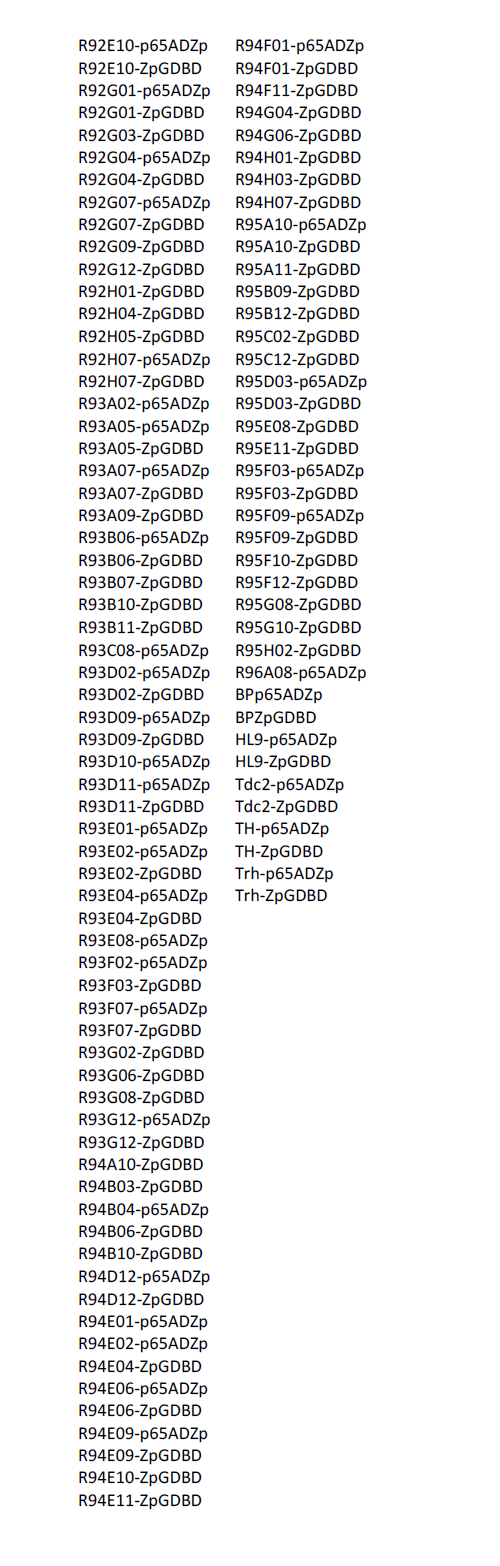

### Using hemidriver lines to generate split-GAL4 lines with desired expression patterns

The first step in generating a split-GAL4 line for a given cell type is screening the images of generation 1 GAL4 (Jenett *et al*. 2012; www.janelia.org/gal4-gen1) to identify enhancers that drive expression in the desired cell type. Once a list of enhancers has been identified, AD and DBD hemidrivers that use those enhancers can be selected from the lines in Table 1 and crossed using a cross scheme such as that shown in Figure 1.

**Figure 1.**
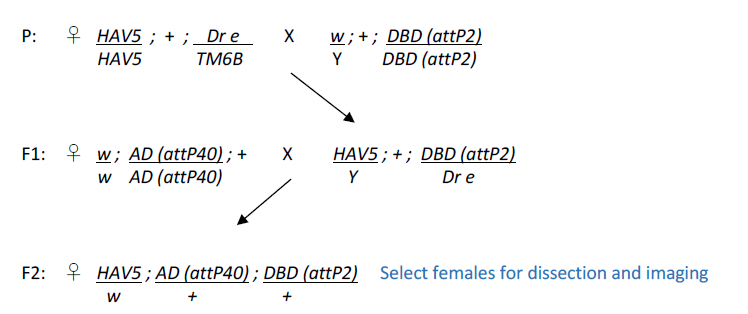
Genetic crosses used for intersectional screening of Split-GAL4s. The genotype of the HAV5 reporter line is *pJFRC200-10XUAS-IVS-myr::smGFP-HA in attP18, pJFRC216-13XLexAop2-IVS-myr::smGFP-V5 in su(Hw)attP8;; Dr e/TM6B*.

The F2 generation of potentially useful AD and DBD hemidriver combinations is then screened for GAL4-driven expression. Typically between 5 and 30 percent of the intersections show expression in a restricted set of cells. However, only about 5 percent of the attempted intersections generated flies that lack detectable expression in off-target cells and could be considered specific for the target cell type. How much off-target expression is acceptable depends on the intended use of the lines. A few off-target cells may not be a problem if the intended use of the split-GAL4 line is anatomical analysis, functional imaging, or as a guide for the placement of a recording electrode. On the other hand, for behavioral screens involving activation of cell types, and to a lesser extent inactivation of cell types, off-target expression can be confounding; such off-target expression can also occur in non-neuronal cells such as muscle, which will not be observed if only dissected nervous systems are examined (see Aso *et al*. 2014). Having multiple independent intersections for a given cell type can be helpful here, as these lines will almost always have different off-target expression. Use of multiple split-GAL4 lines for a cell type that were derived using different hemidriver lines also decreases the chance of being misled by effects of genetic background; while the general background of all the lines is the same there may be mutations associated with the chromosomes carrying the individual AD or DBD transgenes. These could affect fly behavior but are less likely to be the same for multiple AD/DBD combinations. Making the lines more specific by “triple intersections” (see for example, Dolan *et al*. 2017) can also be attempted. Finally, it is important to keep in mind that the expression pattern observed with any GAL4 driver depends on the chromosomal location and number of UASs carried by the indicator gene (see Pfeiffer *et al.* 2010 and Aso *et al.* 2014 for examples). For this reason, it is best to use indicator and effector lines inserted at the same site; in the ideal case, both the indicator and effector functions are provided by the same gene product, for example, 20XUAS-IVS-CsChrimson-mVenus (Klapoetke *et al.* 2014).

Screening is rapid enough that this has generally not been a limiting factor; however, several factors significantly influence the screening success rate:

1. The degree of certainty with which the selected enhancers can be said to express in the desired cell-type is perhaps the largest factor affecting success. Many generation 1 GAL4 lines express in patterns that are too broad to allow identification of individual cell types with confidence and different cell types may have indistinguishable morphologies at this level of imaging resolution. In some cases, we have found it helpful to use stochastic labeling with the Multi-Color-Flip-Out (MCFO) method (Nern *et al*. 2015) to visualize the morphologies of individual cells in generation 1 lines as a way of discerning what cell types are present.
2. Because expression patterns vary with insertion site, the hemidriver expression pattern may differ from that of the generation 1 GAL4 line with the same enhancer because the AD hemidrivers are inserted into attP40 and the generation 1 lines are inserted in attP2. Different insertion sites for the AD and DBD hemidrivers were used to avoid transvection (Mellert and Truman, 2012) and to make it possible to generate stable split-GAL4 lines with both an AD and DBD hemidriver. For many of the enhancers used in Table 1, enhancer-LexA constructs inserted in attP40 have been generated and imaged (www.janelia.org/gal4-gen1) and these images can serve as a useful guide for the expression pattern of the enhancer inserted at that landing site. In cases where the expression pattern is not reproduced in attP40 and the number of enhancers for the desired cell type is limited, we have had success by re-injecting the AD hemidriver into additional landing sites that were selected to be more “similar” to attP2, such as JK22C and JK73A (Knapp *et al*. 2015). Since the DBD hemidrivers are inserted into the same site as the generation 1 GAL4 lines, they reliably reproduce the GAL4 expression pattern. For this reason, a useful strategy is to use an AD hemidriver that is empirically known to express in the target cell type and cross it to DBD hemidrivers from a set of lines selected based on anatomy.
3. A successful intersection requires the identification of at least two generation 1 lines that express in the cell type of interest; for some cell types, the lines in Table 1 will be insufficient for this purpose. In this case there is little choice but to expand the set of hemidriver lines. For example, we have been able to obtain successful intersection in such cases by also incorporating the set of hemidrivers described in the accompanying paper (Tirian *et al*., 2017) that were generated in the same vectors, but used a different set of enhancers. Also, additional hemidriver constructs can be generated if an enhancer is identified for which a hemidriver line does not already exist in either collection. We note that the set of hemidriver lines in Table 1 uses less than half the enhancers described in Jenett *et al*. (2102) and the selection of these enhancers was biased by the anatomical interests of a small number of laboratories. The hemidrivers in these collections can also be used in combinations with hemidrivers generated by other strategies (Gohl *et al*. 2011, Diao *et al*. 2015, Simpson 2016); for example, Aso *et al*. (2014) used a hemidriver made using a large chromosomal region of the tyrosine hydroxylase (TH) gene to make split-GAL4 lines for subsets of dopaminergic neurons.

Once combinations of hemidrivers that yield successful intersections have been identified in the screening process, we generally construct stable split-GAL4 lines that carry a combination of the two hemidrivers, using the cross scheme shown in Figure 2. In cases where uniformity of genetic background is of paramount importance, the hemidrivers can be backcrossed to a common stock before making stable lines. This is usually not an issue, however, as in the general use case, the split-GAL4 line’s chromosomes are heterozygous with those of a common indicator or effector line in the experimental flies that are being assayed.

**Figure 2.**
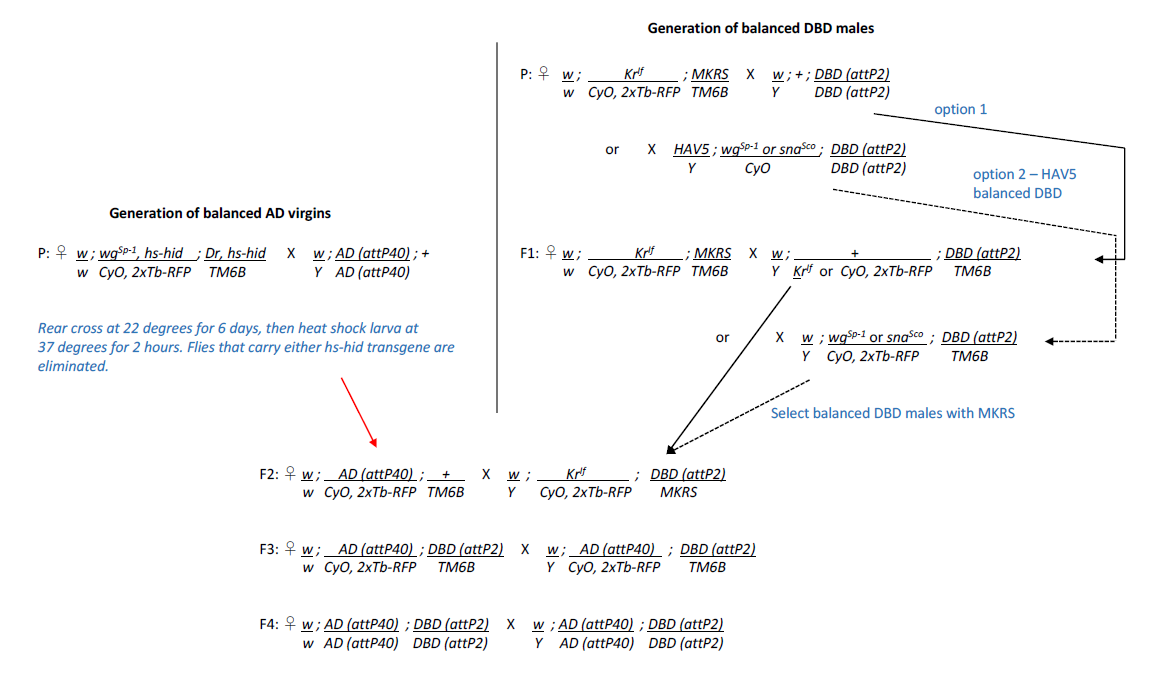
Genetic cross scheme to create stable split-GAL4 combinations.

## Concluding Remarks

Using the combined hemidriver collections described in this paper and Tirian *et al*. (2017), our experience in several brain areas suggests that it will be possible to obtain useful split-GAL4 lines for more than three quarters of the cell types in the adult fly brain. By making additional hemidrivers this percentage could likely be increased. However, we have also found cases where it has not been possible to make split-GAL4 lines that separate two closely related, but clearly morphologically distinct, cell types. This may be a fundamental limitation based on the biology of combinatorial gene control; there may simply be no two enhancers that uniquely share an intersection in those individual cell types.

We anticipate rapid progress on electron microscopic level connectomics in *Drosophila* (for example, Zheng *et al*. 2017; Takemura *et al*. 2017; Eichler *et al*. 2017). These efforts will produce the morphologies of all the neurons participating in particular circuits. Such cell shapes can serve as templates for searching for enhancers that drive expression in the same cell types. These enhancers can then be used to generate the split-GAL4 drivers needed to manipulate those cells or image their activity. Clean drivers, as can be generated by these methods, will permit the manipulation of specific cell types in freely moving flies. Generating these lines will be, of necessity, a community effort, accomplished most efficiently by using shared reagents such as those described here.

## Acknowledgements

We thank Don Hall, Dona Fetter, James McMahon, Danielle Ruiz, Pria Chang, Ling-Yu Liu, Monti Mercer, Grace Zheng, Scarlett Harrison, Guillermo Gonzalez, Brandi Sharp, Tam Dang, Todd R. Laverty for help with fly husbandry and stock generation and Chris Murphy for molecular biology assistance with making the hemidriver constructs. We thank Yoshi Aso, Michael-John Dolan, Arnim Jenett, Aljoscha Nern, Jim Truman, Tanya Wolff and other users of the collection for helping to select the enhancers used for hemidriver generation and for sharing their experiences in using the lines to generate split-GAL4 lines.

## References

Aso Y, Hattori D, Yu Y, Johnston RM, Iyer NA, Ngo TT, Dionne H, Abbott LF, Axel R, Tanimoto H, Rubin GM. The neuronal architecture of the mushroom body provides a logic for associative learning. eLife. 2014 Dec 23;3:e04577. doi:10.7554/eLife.04577.

Diao F, Ironfield H, Luan H, Diao F, Shropshire WC, Ewer J, Marr E, Potter CJ, Landgraf M, White BH. Plug-and-Play Genetic Access to Drosophila Cell Types Using Exchangeable Exon Cassettes Cell Rep. 2015 Mar 3; 10(8): 1410–1421. doi:10.1016/j.celrep.2015.01.059.

Dolan MJ, Luan H, Shropshire WC, Sutcliffe B, Cocanougher B, Scott RL, Frechter S, Zlatic M, Jefferis GSXE, White BH. Facilitating neuron-specific genetic manipulations in Drosophila melanogaster using a split GAL4 repressor. Genetics. 2017 Jun;206(2):775–784. doi:10.1534/genetics.116.199687.

Eichler K, Li F, Litwin-Kumar A, Park Y, Andrade I, Schneider-Mizell CM, Saumweber T, Huser A, Eschbach C, Gerber B, Fetter RD, Truman JW, Priebe CE, Abbott LF, Thum AS, Zlatic M, Cardona A. The complete connectome of a learning and memory centre in an insect brain. Nature. 2017 Aug 9;548(7666):175–182. doi: 10.1038/nature23455.

Gohl DM, Silies MA, Gao XJ, Bhalerao S, Luongo FJ, Lin CC, Potter CJ, Clandinin TR. A versatile in vivo system for directed dissection of gene expression patterns. Nat Methods. 2011 Mar;8(3):231–7. doi: 10.1038/nmeth.1561.

Groth AC, Fish M, Nusse R, Calos MP. Construction of transgenic Drosophila by using the site-specific integrase from phage phiC31. Genetics. 2004 Apr;166(4):1775–82. doi: 10.1534/genetics.166.4.1775.

Jenett A, Rubin GM, Ngo TT, Shepherd D, Murphy C, Dionne H, Pfeiffer BD, Cavallaro A, Hall D, Jeter J, Iyer N, Fetter D, Hausenfluck JH, Peng H, Trautman ET, Svirskas RR, Myers EW, Iwinski ZR, Aso Y, DePasquale GM, Enos A, Hulamm P, Lam SC, Li HH, Laverty TR, Long F, Qu L, Murphy SD, Rokicki K, Safford T, Shaw K, Simpson JH, Sowell A, Tae S, Yu Y, Zugates CT. A GAL4-driver line resource for Drosophila neurobiology. Cell Rep. 2012 Oct 25;2(4):991–1001. doi: 10.1016/j.celrep.2012.09.011.

Knapp JM, Chung P, Simpson JH. Generating customized transgene landing sites and multi-transgene arrays in Drosophila using phiC31 integrase. Genetics. 2015 Apr;199(4):919–34. doi: 10.1534/genetics.114.173187.

Klapoetke NC, Murata Y, Kim SS, Pulver SR, Birdsey-Benson A, Cho YK, Morimoto TK, Chuong AS, Carpenter EJ, Tian Z, Wang J, Xie Y, Yan Z, Zhang Y, Chow BY, Surek B, Melkonian M, Jayaraman V, Constantine-Paton M, Wong GK, Boyden ES. Independent optical excitation of distinct neural populations. Nat Methods. 2014 Mar;11(3):338–46. doi: 10.1038/nmeth.2836.

Luan H, Peabody NC, Vinson CR, White BH. Refined spatial manipulation of neuronal function by combinatorial restriction of transgene expression. Neuron. 2006 Nov 9;52(3):425–36. doi: 10.1016/j.neuron.2006.08.028.

Markstein M, Pitsouli C, Villalta C, Celniker SE, Perrimon N. Exploiting position effects and the gypsy retrovirus insulator to engineer precisely expressed transgenes. Nat Genet. 2008 Apr;40(4):476–83. doi: 10.1038/ng.101.

Mellert DJ, Truman JW. Transvection is common throughout the Drosophila genome. Genetics. 2012 Aug;191(4):1129–41. doi: 10.1534/genetics.112.140475.

Nern A, Pfeiffer BD, Rubin GM. Optimized tools for multicolor stochastic labeling reveal diverse stereotyped cell arrangements in the fly visual system. Proc Natl Acad Sci U S A. 2015 Jun 2;112(22):E2967–76. doi: 10.1073/pnas.1506763112.

Pfeiffer BD, Jenett A, Hammonds AS, Ngo TT, Misra S, Murphy C, Scully A, Carlson JW, Wan KH, Laverty TR, Mungall C, Svirskas R, Kadonaga JT, Doe CQ, Eisen MB, Celniker SE, Rubin GM. Tools for neuroanatomy and neurogenetics in Drosophila. Proc Natl Acad Sci U S A. 2008 Jul 15;105(28):9715–20. doi: 10.1073/pnas.0803697105.

Pfeiffer BD, Ngo TT, Hibbard KL, Murphy C, Jenett A, Truman JW, Rubin GM. Refinement of tools for targeted gene expression in Drosophila. Genetics. 2010 Oct;186(2):735–55. doi: 10.1534/genetics.110.119917.

Simpson JH. Rationally subdividing the fly nervous system with versatile expression reagents. J Neurogenet. 2016 Sep - Dec;30(3-4):185–194. doi: 10.1080/01677063.2016.1248761.

Takemura SY, Aso Y, Hige T, Wong A, Lu Z, Xu CS, Rivlin PK, Hess H, Zhao T, Parag T, Berg S, Huang G, Katz W, Olbris DJ, Plaza S, Umayam L, Aniceto R, Chang LA, Lauchie S, Ogundeyi O, Ordish C, Shinomiya A, Sigmund C, Takemura S, Tran J, Turner GC, Rubin GM, Scheffer LK. A connectome of a learning and memory center in the adult Drosophila brain. eLife. 2017 Jul 18;6. pii: e26975. doi: 10.7554/eLife.26975.

Tirian L, Fellner M, Dickson BJ. The VT GAL4, LexA, and split-GAL4 collection for targeted expression in the Drosophila nervous system (bioRxiv).

Tuthill JC, Nern A, Holtz SL, Rubin GM, Reiser MB. Contributions of the 12 neuron classes in the fly lamina to motion vision. Neuron. 2013 Jul 10;79(1):128–40. doi: 10.1016/j.neuron.2013.05.024.

Wu M, Nern A, Williamson WR, Morimoto MM, Reiser MB, Card GM, Rubin GM. Visual projection neurons in the Drosophila lobula link feature detection to distinct behavioral programs. eLife. 2016 Dec 28;5. pii: e21022. doi: 10.7554/eLife.21022.

Zheng Z, Lauritzen JS, Perlman, Robinson CG, Nichols M, Milkie D, Torrens O, Price J, Fisher CB, Sharifi N, Calle-Schuler SA, Kmecova L, Ali IJ, Karsh B, Trautman ET, Bogovic J, Hanslovsky P, Jefferis GSXE, Kazhdan M, Khairy K, Saalfeld S, Fetter RD, Bock DD. A complete electron microscopy volume of the brain of adult Drosophila melanogaster. doi: https://doi.org/10.1101/140905. http://biorxiv.org/content/early/2017/05/22/140905

